# Aggregating Residue-Level Protein Language Model Embeddings with Optimal Transport

**DOI:** 10.1101/2024.01.29.577794

**Authors:** Navid NaderiAlizadeh, Rohit Singh

## Abstract

Protein language models (PLMs) have emerged as powerful approaches for mapping protein sequences into embeddings suitable for various applications. As protein representation schemes, PLMs generate per-token (i.e., per-residue) representations, resulting in variable-sized outputs based on protein length. This variability poses a challenge for protein-level prediction tasks that require uniform-sized embeddings for consistent analysis across different proteins. Previous work has typically used average pooling to summarize token-level PLM outputs, but it is unclear whether this method effectively prioritizes the relevant information across token-level representations. We introduce a novel method utilizing optimal transport to convert variable-length PLM outputs into fixed-length representations. We conceptualize per-token PLM outputs as samples from a probabilistic distribution and employ sliced-Wasserstein distances to map these samples against a reference set, creating a Euclidean embedding in the output space. The resulting embedding is agnostic to the length of the input and represents the entire protein. We demonstrate the superiority of our method over average pooling for several downstream prediction tasks, particularly with constrained PLM sizes, enabling smaller-scale PLMs to match or exceed the performance of average-pooled larger-scale PLMs. Our aggregation scheme is especially effective for longer protein sequences by capturing essential information that might be lost through average pooling.

## Introduction

Understanding the sequence–structure–function relationship for proteins is one of the grand challenges of biology. Among the problems defined around these relationships, a prominent class of problems comprises tasks where some structural or functional property of a protein is to be predicted from its sequence. The possible set of prediction tasks in this class is very diverse, including both classification (e.g., “is the protein a kinase?”) and regression (e.g., the melting temperature of the protein), as well as tasks involving auxiliary inputs (e.g., small molecule SMILES representations, for predicting drug-target interactions). Any such problem can be formulated as a prediction task over a set of proteins, where the input consists of a protein sequence and the output (a label or number) is at the level of the *entire protein*, rather than its constituent amino acids. Any model that addresses such a problem formulation will necessarily have one or more steps where information across the constituent amino acids of the protein is summarized into a protein-level estimate.

The problem of appropriately aggregating amino acid-level information has become particularly pressing with the advent of protein language models (PLMs). PLMs, which are trained on massive corpora of protein sequence data using self-supervised learning, are able to build internal representations that capture evolutionary constraints on protein sequences. Since these representations comprehensively capture constraints on protein function and structure, PLMs have proven powerful in a wide range of tasks, from structure prediction to interaction prediction and protein design. For many protein sequence-based property-prediction tasks, PLM-based approaches are now the state-of-the-art [25; 18; 39]. Given a sequence of length *n*, a pre-trained PLM produces an embedding of dimensionality ℝ^*n×d*^, where *d* is the per-amino acid (i.e., per-token) embedding dimensionality. The token-specific embedding captures not only biochemical information about the token (i.e., the amino acid) but also the local and global properties of the protein.

To aggregate these per-token embeddings into a protein-level representation, the most common approach has been to simply “average pool:” take the mean of each feature dimension along the length of the protein to produce an embedding in ℝ^*d*^ [4; 46; 47; 39; 40]. While other pooling approaches, such as “max pooling” (taking the maximum of the set) [12] or “softmax pooling” (i.e., average pooling after exponentiation, then logarithmized) [38] are also sometimes used, average pooling is typically preferred for convenience, speed, and simplicity. However, it weighs each amino acid’s representation equally. This is unrealistic—often, there are specific residues in the protein that are particularly important (e.g., the residues at an active site). Even when the per-token PLM representation does contain information distinguishing such residues from others, these distinctions might be lost during average pooling, which could be especially problematic for longer protein sequences. We note that such considerations are not limited to PLM-based embeddings: in many task-specific neural network architectures, e.g., PIPR for PPI prediction [8], average pooling is used to summarize variable-length intermediate representations.

In this work, we present a novel approach to aggregate variable-length protein representations. Like average pooling, max pooling, and related approaches, our method is permutation invariant: it considers the per-token embeddings as a *set* rather than a *sequence*, under the assumption that the PLM backbone fully embeds the sequential properties of the protein data in its output residue-level representations. Let **p**_*j*_ be a protein of length *n*_*j*_, so that its PLM embedding can be represented as 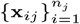, with each **x**_*ij*_ ∈ ℝ^*d*^. Our work seeks to learn a set of *m* reference embeddings 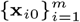, with **x**_*i*0_ ∈ ℝ^*d*^, that can characterize any variable-length representation. Conceptually, this is analogous to learning a task-specific basis representation in ℝ^*d*^. The intuition underpinning our work is to think of 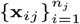 and 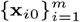 as empirical probability distributions in ℝ^*d*^ and to formulate 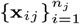’s distance from the reference set 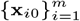 as an optimal transport calculation.

Our work builds upon optimal transport (OT) based approaches in computer vision to characterize sets of observations. We borrow from previous advances to deploy the OT intuition effectively and in a scalable fashion. In particular, we use “slices” in the embedding space, learnable directions in ℝ^*d*^ onto which the input and reference token-sets 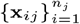 and 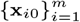 are projected. These projections correspond to 1-D probability distributions. On such distributions, OT distances can be computed efficiently, and an ensemble of *L* slices serves to efficiently characterize the separation between input and reference sets.

The key conceptual advance of our work is unlocking task-specific learnability as a key component of PLM-based machine learning models. Broadly, these models can be thought of as pipelines of three segments: an initial *sequence* segment, a *summarization* segment, and a final *prediction* segment. For example, the sequence segment may consist of transformer layers, while the prediction segment may consist of a feed-forward network. However, if average pooling is used for summarization, no task-specific learning can happen there and will need to happen only in the sequence or the prediction segment. With our innovation, the summarization segment also becomes learnable, offering greater flexibility in matching the architecture of the neural network to the biological intuitions underlying the task.

Our work also has the potential to introduce interpretability into systems that would otherwise be opaque. The set of *m* reference embeddings learned by the system can serve as useful archetypes for the task at hand. For instance, in the computer vision context, it was shown that simply being able to associate the variable-length representation with one of the references can be informative about the typicality of the underlying object.

We apply our sliced-Wasserstein embedding (SWE) approach to four protein property prediction tasks: binary drug-target interaction prediction, out-of-domain drug-target affinity prediction, subcellular localization, and enzyme commission (EC) prediction. SWE broadly outperforms other pooling mechanisms, including average pooling, especially on small and moderate-sized PLMs. Notably, the onerous GPU memory requirements of the largest PLM architectures suggest that the performance boost of SWE could be critical to democratizing access to PLMs for researchers with limited GPU resources. SWE especially shines over average pooling for longer protein sequences, hence confirming our hypothesis that average pooling results in a more substantial loss of information when dealing with larger numbers of token-level representations.

## Related Work

### Protein Language Models

Large language models (LLMs) have become the predominant tools for modeling sequential natural language data. The success of LLMs, which mostly rely on attention-based transformer architectures [48], has inspired researchers working with biological data to use similar ideas for analyzing protein sequences. In particular, the availability of massive protein sequence datasets has given rise to large-scale protein language models (PLMs), such as ESM [36; 24], ProtBert [13], ProtGPT2 [14], SaProt [43], xTrimoPGLM [7], and DPLM [52], to name a few. These models are mostly trained using unlabeled protein sequence data in a self-supervised way, where the goal is to train the model to predict a token that has been replaced with a special *mask* token using its surrounding context, i.e., other amino acids in the sequence. Such masking-based unsupervised training leads to token-level representations that have been shown to provide state-of-the-art performance in a wide array of downstream tasks, such as protein folding [50], variant effect prediction [6], peptide generation [9], antibody design [53], and prokaryotic gene prediction [45].

The transformer architectures in LLMs, in general, and PLMs, in particular, produce residue-level representations that need to be summarized and aggregated for protein-level downstream tasks since different amino acid sequences have varying lengths. The question of aggregating a set of elements into a fixed-length representation is the key behind the research on *set representation learning*, which we discuss next.

### Set Representation Learning

The goal of set representation learning is to map an unordered collection of elements into an embedding that is invariant to the permutation of the set elements and whose size is independent of the input set size [35; 51]. Deep Sets [54] and Janossy Pooling [30] are two seminal studies in this area, where the set embedding is modeled as a function of the sum or average of permutation-sensitive functions applied to all elements or all permutations of the input set. Follow-up work has leveraged ideas based on transformers [22] and optimal transport [20; 32; 31; 28], among others, for deep learning on sets, and demonstrated their efficacy in a variety of learning settings, including point cloud classification [34], graph representation [29], and multi-agent reinforcement learning [44].

In the context of PLMs, prior work has, for the most part, used *average pooling* to summarize the token-level embeddings into universal protein-level embeddings [4; 46; 47; 39; 40], and past research on other pooling methods is scarce [41; 42; 17; 12]. Averaging is a simple, unparameterized, permutation-invariant, and size-invariant function, which warrants its selection as the most natural and intuitive choice for aggregating PLM outputs. Nevertheless, it is unknown whether other, more sophisticated aggregation mechanisms could unlock additional performance gains compared to average pooling when used in conjunction with state-of-the-art PLMs. In this paper, we give an affirmative answer to this question by proposing a parameterized aggregation operation based on ideas from optimal transport to summarize residue-level embeddings generated by pre-trained PLMs into fixed-length protein-level embeddings.

## Methods

### Problem Formulation

Consider a protein’s primary amino acid sequence of length*n* ∈ *ℕ*, denoted by **p** = (*p*_1_, …, *p*_*n*_) ∈ *𝒫*^*n*^, where ℕ denotes the set of natural numbers, i.e., positive integers, 𝒫 represents the residue alphabet. We gather the set of all possible protein sequences of arbitrary lengths into a set

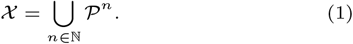

The goal of protein-level representation learning is to find a function *ψ*(*·*; *θ*_*ψ*_) : 𝒳 → *ℝ*^*d*^, parameterized by a finite-dimensional set of parameters *θ*_*ψ*_ ∈ Θ_*ψ*_ (see Figure 1-a). Observe that the dimensionality of the embedding space, i.e., *d*, and the size of the model parameter space, i.e., |Θ|, are *independent* of the length of the input protein sequence. In other words, the function *ψ*(*·*; *θ*_*ψ*_) should be able to map any given protein sequence of arbitrary length to a fixed-size representation in ℝ^*d*^.

**Fig. 1.**
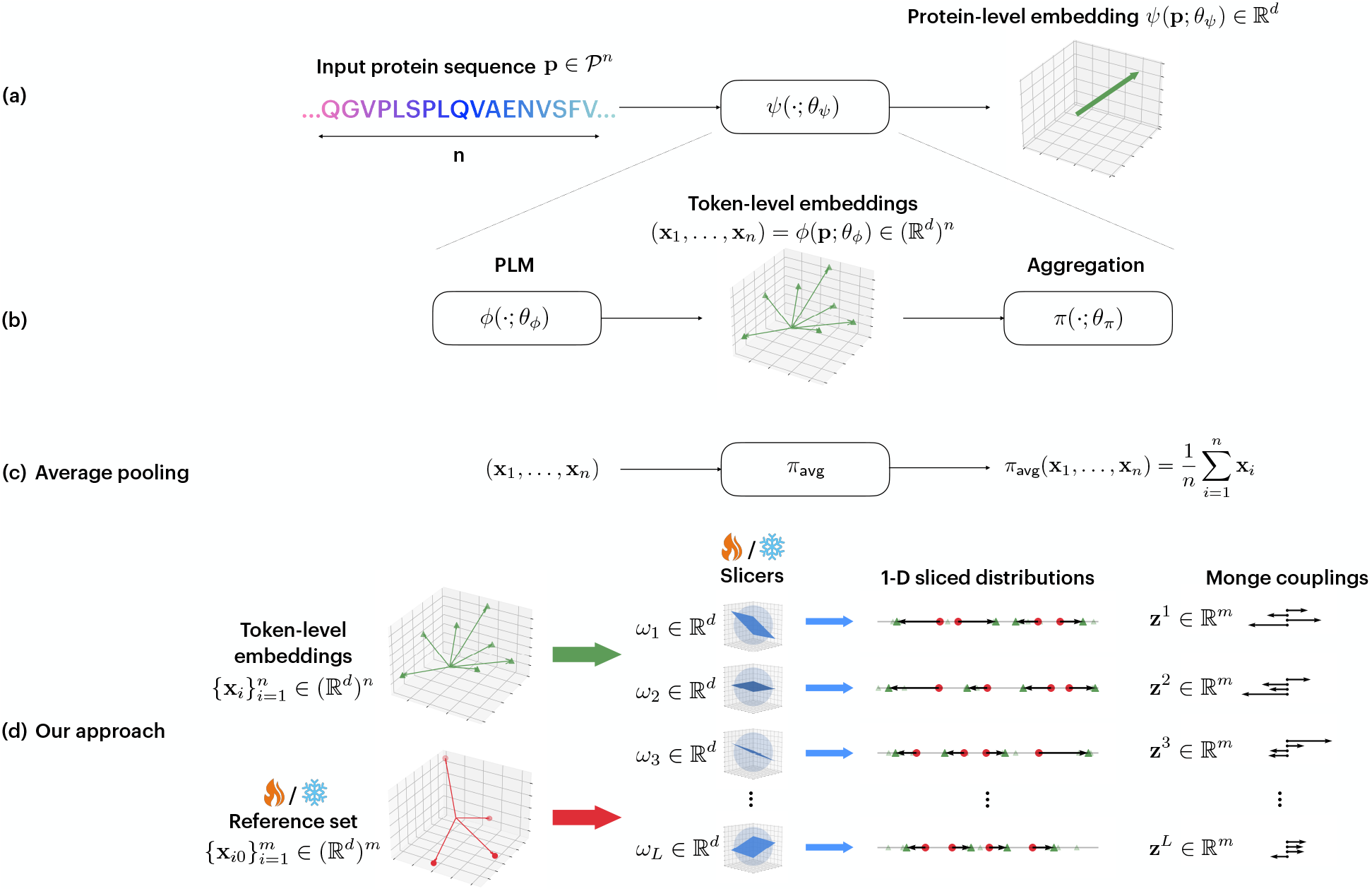
Overview of the proposed method and comparison with average pooling. (a) We consider a parameterized protein representation learning function *ψ*(*·*; *θ*_*ψ*_), which takes as input a protein’s amino acid sequence of arbitrary length and produces a fixed-length protein-level embedding at its output. (b) Breaking down the representation learning pipeline, we first pass the amino acid sequence through a pre-trained protein language model (PLM) *ϕ*(*·, θ*_*ϕ*_), thereby generating a set of token-level embeddings (**x**_1_, …, **x**_*n*_), each residing in ℝ^*d*^. An aggregation function *π*(*·, θ*_*π*_) subsequently summarizes these embeddings into a protein-level embedding, whose size does not depend on the sequence length *n*. (c) Average pooling, which is most commonly used in the literature, simply takes the mean of the residue-level embeddings to derive the protein-level embedding. (d) Our proposed sliced-Wasserstein embedding aggregation module, which is based on comparing the probability distribution underlying the token-level embeddings, and a trainable reference probability distribution. Since such comparison is non-trivial in a high-dimensional space, we pass the token-level embeddings and a set of *m* reference elements through a set of *L* trainable linear slicing operations 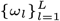, which map the embeddings and reference elements into *L* pairs of 1-dimensional distributions. The Monge couplings between these 1-D distributions are then calculated based on sorting and interpolation operations, and subsequently combined to generate the final fixed-length protein-level embedding. Note that in the absence of enough downstream training data, the parameters of the reference set and slicers can be left frozen, and only the final *m*-dimensional combination of the Monge couplings need to be learned; we refer to this variant of our proposed pooling method as “SWE_Simple.”

Assuming the availability of a (labeled) protein sequence dataset 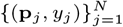, where **p**_*j*_ ∈ *𝒳, y*_*j*_ ∈ *𝒴*, ∀*j* ∈ {1, …, *N*}, with 𝒴 denoting the set of all possible labels, the parameters of the representation function *ψ* are typically derived via an empirical risk minimization (ERM) problem,

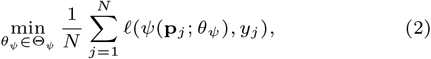

where 𝓁 : ℝ^*d*^ *×𝒴* → *ℝ* denotes a loss function. While resembling a supervised learning setting, note that the formulation in also includes an unsupervised learning scenario, where the labels *y*_*j*_ are trivial/non-existent and the loss function 𝓁 does not depend on *y*.

A common initial building block used to parameterize the representation function *ψ* is a protein language model (PLM). We denote a PLM by a parameterized function *ϕ*(*·, θ*_*ϕ*_) : 𝒫^*n*^ → (ℝ^*d*^)^*n*^, ∀*n* ∈ *ℕ*. This implies that a given PLM maps each amino acid in an input protein sequence into an individual embedding in ℝ^*d*^. The parameters of the PLM, i.e., *θ*_*ϕ*_, are usually trained by a masking-based objective, where some amino acid identities in the input sequences are masked by random tokens, and the model is trained to predict the correct amino acids using the corresponding output representations.

In this paper, our goal is on the aggregation, or pooling, function that bridges the gap between PLM-generated token-level outputs and the final protein-level representation. More formally, we are interested in an informative aggregation function *π*(*·*; *θ*_*π*_) : (ℝ^*d*^)^*n*^ → ℝ, ∀*n* ∈ *ℕ*, parameterized by *θ*_*π*_ ∈ Θ_*π*_. Taken together, the composition of the PLM *ϕ* and the aggregation function *π* constitutes the end-to-end protein-level representation learning pipeline (see Figure 1-b); for any given protein sequence **p** ∈ *𝒳*, its representation is derived as

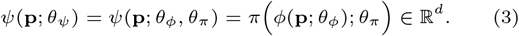

We assume that the PLM is pre-trained and its parameters, *θ*_*ϕ*_, are frozen. Therefore, we are primarily interested in training the aggregation function *π*(*·*; *θ*_*π*_) for downstream prediction tasks.

### Proposed Pooling Method

A simple and prominent example of an aggregation function is the *averaging* operation, i.e., 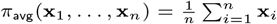, which is indeed unparameterized (i.e., Θ_*ψ*_ = ∅; see Figure 1-c). While used extensively in prior work in the protein representation learning literature (see, e.g., [40; 39]), average pooling may not capture the entirety of the information that is present in the per-token embeddings. This calls for methods that are able to capture such information in the aggregated representations.

In this paper, we propose to use sliced-Wasserstein distances from optimal transport [11; 21], and in particular, *sliced-Wasserstein embedding* (SWE) [31; 28] to aggregate the token-level representations into a universal protein-level presentation. The main idea behind SWE is to treat the token-level embeddings as samples drawn from an underlying probability distribution, and then find the optimal transportation plan that maps that distribution to a trainable reference distribution (see Figure 1-d).

More formally, let 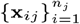 denote the token-level embeddings of the *j*^th^ protein sequence in the dataset, **p**_*j*_, *j* ∈ {1, …, *N*}, of length *n*_*j*_, which are produced by the pre-trained PLM; i.e., 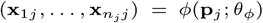. We assume that the embeddings 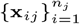 are samples drawn from an underlying distribution 𝔇_*j*_ supported on ℝ^*d*^. We also consider a trainable reference set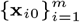 of *m* points in ℝ^*d*^ that are drawn from a reference distribution 𝔇_0_. Since there is no closed-form solution for calculating the optimal Monge coupling [49] between high-dimensional distributions in ℝ^*d*^, we resort to *slicing* operations in order to map these distributions to several one-dimensional distributions, for which the Monge coupling has a closed-form solution. In particular, we consider trainable linear maps 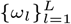, with *ω*_*l*_ ∈ ℝ^*d*^, ∀*l* ∈ {1, …, *L*}, through which each set of token-level embeddings 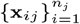, *j* ∈ {0, …, *N*} (including the reference embeddings with *n*_0_ = *m*) is mapped to *L slices*, i.e., sets of one-dimensional points 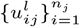, *l* ∈ {1, …, *L*}, where

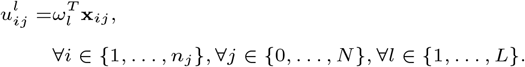

Consider the *l*^th^ slice of a set of token-level embeddings, i.e., 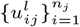 and the corresponding slice of the reference set, i.e., 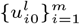. These sets correspond to empirical 1-D distributions 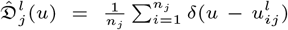 and 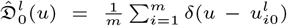, respectively. The Monge coupling between 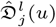 and 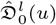 has a closed-form solution, and in particular is an *m*-dimensional vector 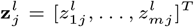 which relies solely on sorting and interpolation operations. In particular, depending on whether the length of the input protein sequence and the size of the reference set are identical, there are two cases:

- **Case 1 (***n*_*j*_ = *m*): In this scenario, the Monge map is simply the difference between the sorted sequences of points in the input slice and the reference slice. More precisely, let *ρ*[*·*] denote the permutation indices obtained by sorting 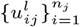. Moreover, let 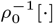 denote the ordering that permutes the sorted set back to the original ordering based on sorting of elements in the reference set 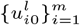. Then, the Monge map elements are given by

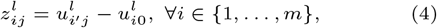

where 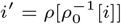.
- **Case 2 (***n*_*j*_ *≠*= *m*): The embedding procedure for this case follows similar steps as in Case 1, with the addition of an interpolation operation. Specifically, we derive the interpolated inverse cumulative distribution function (CDF) of the sliced token-level values, which we denote by 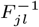. This involves sorting 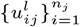, calculating the cumulative sum, and calculating the inverse through interpolation. The Monge coupling elements can then be derived as

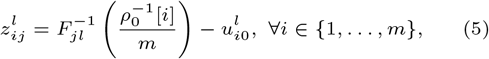

where 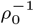 is defined as in Case 1. Observe that (5) reduces to (4) if *n*_*j*_ = *m*.

Once the Monge couplings 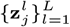 have been derived for all slices, we concatenate them to form the embedding matrix 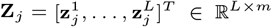. To reduce the dimensionality of the embedding matrix, similarly to [31], we perform a combination across the reference set elements using a learnable projection **w** ∈ ℝ^*m*^ to derive the final protein-level embedding

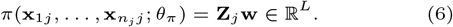

We set *L* = *d*, so that the output embedding dimensionality equals that of average pooling. Besides, to ease the optimization process, we learn the reference elements at the *output* of the slicers, training 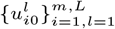 directly instead of 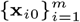.

#### Runtime and Memory Considerations

Note that the trainable SWE parameters include the parameters of the reference set, (ii) the slicers, and (iii) the final combination across reference elements, i.e., 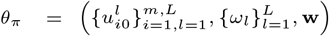. This implies that the total number of parameters in the proposed SWE aggregation function is *mL* + *dL* + *m* = 𝒪 ((*m* + *d*)*L*). We also consider a simpler version of SWE (referred to as “SWE_Simple”), where the reference and slicer parameters are frozen at random, and only the final projection **w** is learned end-to-end, hence leading to only *m* learnable parameters. For the settings we consider in our numerical evaluations, as we describe next, these are negligible overheads as compared to the size of the backbone PLMs.

### Evaluation Settings

We use the state-of-the-art transformer-based ESM-2 [24] and ProGen2 [33] as the PLMs *ϕ*(*·, θ*_*ϕ*_) that generate token-level embeddings. These models have been pre-trained via unsupervised mask-based objectives using tens of millions of unique protein sequences and are shown to encode evolutionary patterns in protein sequences. We evaluate our SWE aggregation method when applied to the outputs of five distinct pre-trained ESM-2 models (with 8M, 35M, 150M, 650M, and 3B parameters), as well as four distinct pre-trained ProGen2 models (small, medium, base, and large).

For the SWE aggregation method, we consider 11 options for the number of reference points, *m* ∈ {100, 200, …, 1000} and two options for freezing the reference and slicer parameters (True/False), leading to 22 different SWE configurations. Note that for the “SWE_Simple” version, the reference and slicer parameters are frozen, hence the validation only happens across the 11 options for the reference set size. In all experiments, we set the number of slicers equal to the output embedding dimensionality of the corresponding PLM, i.e., *L* = *d*.

We evaluate our proposed pooling method across four different downstream tasks as described below. Additional results on protein-protein interaction (PPI) prediction can be found in the Appendix.

- **Drug-Target Interaction (DTI) Prediction**: In this binary classification task, the goal is to predict whether or not a given drug interacts with a target protein. As in [39], for a given drug, we first find its Morgan fingerprint, denoted by **f** ∈ ℝ^*c*^. Using two linear projections **S** ∈ ℝ^*D×c*^ and **V** ∈ ℝ^*D×d*^, we then, respectively, map the drug and the target to a shared co-embedding space ℝ^*D*^, followed by an element-wise non-linearity *σ*(*·*). We then use the cosine similarity between the drug and target embeddings to calculate their interaction probability, which we then optimize using the cross-entropy loss. In particular, for a given training dataset of (drug fingerprint, target, label) triplets 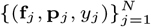, the supervised learning problem in (2) is reformulated as

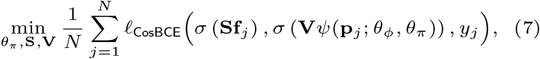

where the optimization occurs over the parameters of the SWE aggregation function, *θ*_*π*_, the drug projector, **S**, and the target projector, **V**, while the parameters of the PLM, *θ*_*ϕ*_, are kept frozen. The loss function 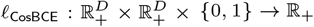 in (7) is defined as

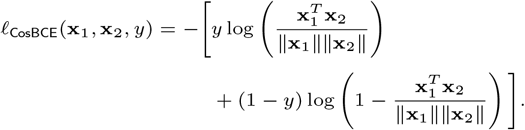 We use `ReLU`(*·*) as the non-linearity *σ*(*·*) to produce both the drug and target embeddings in the non-negative orthant of the embedding space, i.e., 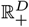. We use two datasets used by [39], namely DAVIS [10] and Binding-DB [26], to evaluate our proposed embedding mechanism in this task.
- **Out-of-Domain Drug-Target Affinity Prediction**: In this regression task, we use the Therapeutics Data Commons (TDC) Drug-Target Interaction Domain Generalization (DTI-DG) Benchmark [16], where the goal is to predict the affinity of the interaction between drugs and protein targets patented between 2019 and 2021 by training on DTI interaction affinities patented in the preceding 5-year window (i.e., 2013–2018). We use the inner product between the drug and target embeddings to predict the interaction affinity and optimize the prediction model parameters using mean squared error (MSE). Especially, we replace the loss function in (7) with 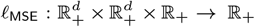, defined as

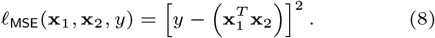
- **Sub-Cellular Localization (SCL)**: This is a 10-class protein property prediction task, where the goal is to predict the localization compartment of a given protein within the cell [1; 42; 23]. Given *N* training samples 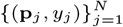, we consider a multi-class classification version of the supervised learning problem (2), i.e.,

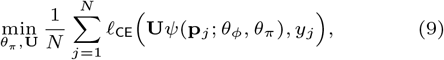

where **U** ∈ ℝ^10*×d*^ denotes a linear classification head, and 𝓁_CE_ : ℝ^10^ *× {*1, …, 10} → ℝ_+_ is defined as

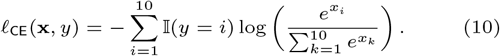
- **Enzyme Commission (EC) Prediction**: We finally consider a binary classification version of the EC number prediction task [15; 55], where we predict whether a given protein is capable of catalyzing biochemical reactions. The formulation of this problem is similar to (9)-(10), where the number of classes is reduced to two, and the classification head is replaced by a two-layer perceptron with a *d*-dimenstional hidden layer and a LeakyReLU non-linearity.

We run our numerical experiments with five different random seeds for 50 epochs. We report the mean and standard deviation of the test performance for the SWE configuration with the highest target validation performance across the 50 training epochs. The target validation performance metric for the binary DTI, DTI-DG, SCL, and EC tasks are set to AUPR (area under the precision-recall curve), PCC (Pearson correlation coefficient), accuracy, and *F*_max_ (*F*_1_-score with the optimal threshold), respectively. More details on the training settings can be found in the Appendix.

While we use average pooling as the main baseline pooling method in all our experiments, for the binary DTI prediction task, we also compare our proposed pooling method with three other baseline pooling methods, including max pooling, *K*-max (average pooling only across top-*K* elements in each dimension) [12], and light attention [42]. Our implementation code can be found at https://github.com/navid-naderi/PLM_SWE.

## Results

### Binary Drug-Target Interaction Prediction

Figure 2 shows the performance of the proposed SWE-based aggregation method for the binary DTI task, evaluated over the DAVIS and Binding-DB datasets. Our proposed SWE aggregation function and its simple variant consistently outperform average pooling in all scenarios and other pooling mechanisms in the majority of the cases. Our method especially shines when i) the PLM is smaller and has lower expressive power (e.g., 8M vs. 3B ESM-2 PLMs or small vs. large ProGen2 PLMs), and ii) there is more data to train the SWE parameters (Binding-DB vs. DAVIS). While the latter point is intuitive since SWE aggregation introduces additional trainable parameters and is, therefore, prone to overfitting, the former point is very significant from an efficiency point of view. In particular, given limited computational resources, where one is constrained to using smaller PLMs, our proposed aggregation operation can, in most cases, push the performance of the smaller PLMs to similar, or even higher, levels than larger PLMs whose outputs are aggregated using averaging. Given the presence of enough training data, especially in the case of Binding-DB, our results demonstrate that there are also gains to be achieved for larger PLM backbones with billions of parameters when using SWE-based aggregation as compared to other pooling methods. However, in the absence of sufficient training data in the case of the DAVIS dataset, SWE mostly falls back to its simple version, where the reference and slicer parameters are randomly initialized and kept frozen. We provide the results on the test AUROC (area under the receiver operating characteristic curve) metric in the Appendix.

**Fig. 2.**
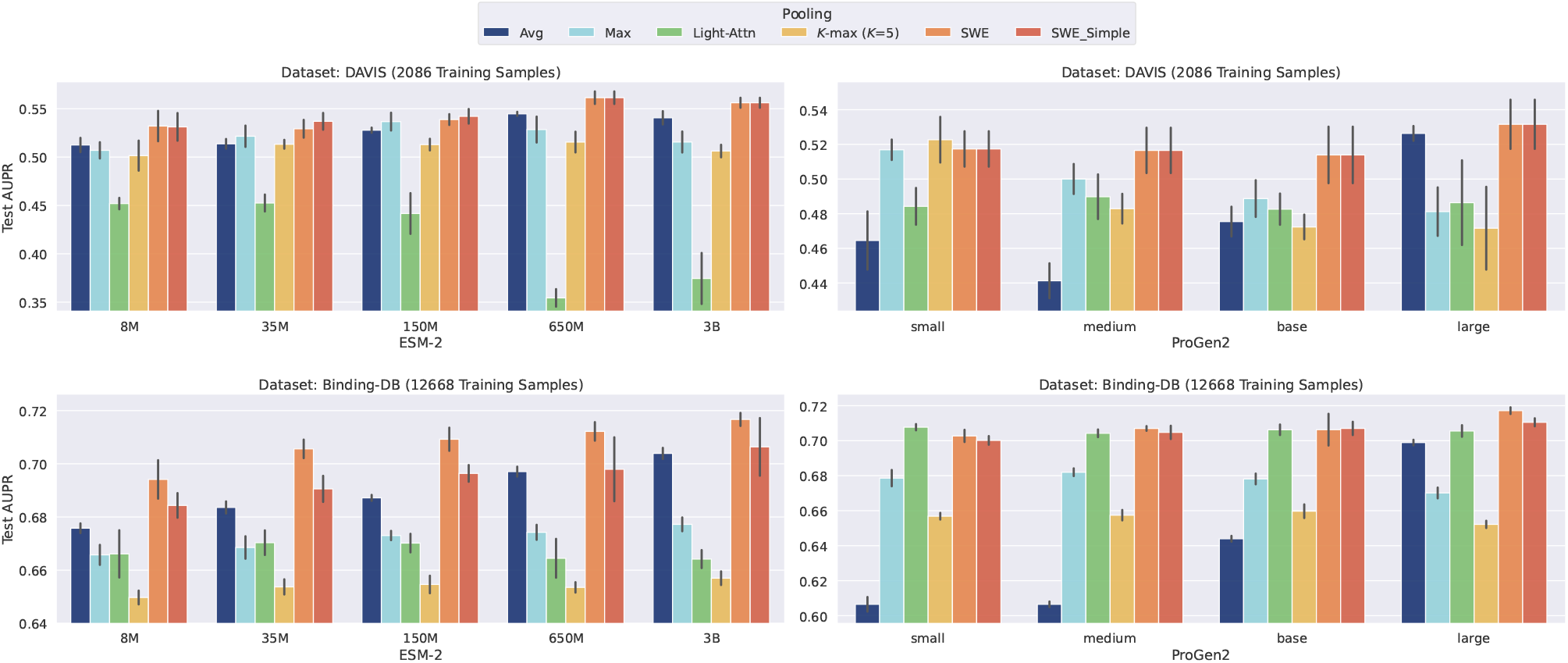
Test AUPR performance of SWE and SWE_Simple as compared to baseline pooling methods for the binary drug-target interaction task on the DAVIS (top) and Binding-DB (bottom) datasets. Performance is evaluated with two families of PLMs, namely ESM-2 (left) and ProGen2 (right).

### Impact of Protein Length on SWE Gains

Figure 3 illustrates how SWE’s gains over average pooling in the Binding-DB DTI prediction task with ProGen2 PLMs correlate with the length of proteins’ amino acid sequences. Quite interestingly, and especially with smaller PLM architectures, the gains are significantly higher for longer protein sequences. This is intuitive since drug-target interactions are typically governed by only a few amino acids, and while such information could get lost when the token-level representations generated by PLMs are simply averaged, our proposed method can extract more meaningful protein-level representations, especially for smaller PLMs and longer sequences. Similar results for the gains in test AUROC can be found in the Appendix.

**Fig. 3.**
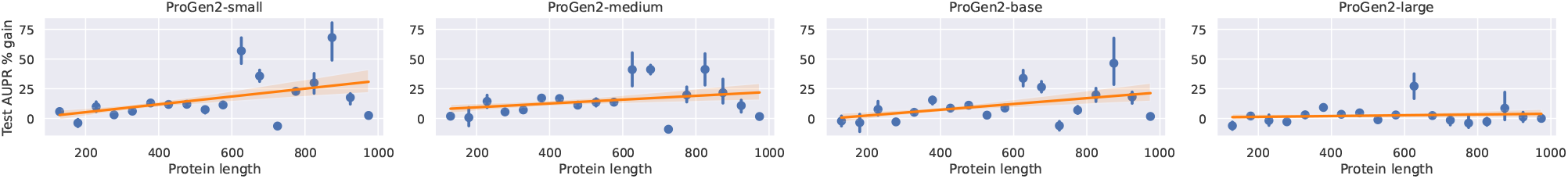
Test AUPR gains of SWE over average pooling vs. the lengths of the proteins’ amino acid sequences across four ProGen2 PLMs on Binding-DB.

### Out-of-Domain Drug-Target Affinity Prediction

Figure 4-a shows the test PCC levels achieved by our proposed SWE aggregation method (and its simple version) as compared to average pooling. We observe that in this task, the SWE pooling mechanism performs better than average pooling for all the four considered ESM-2 PLMs, and interestingly, as in the DAVIS dataset, our hyperparameter grid search prefers to keep the reference and slicer parameters frozen for all PLM sizes. With a test PCC of 0.552 *±* 0.006, our proposed method ranks second in the TDC DTI-DG leaderboard^1^ as of this writing.

**Fig. 4.**
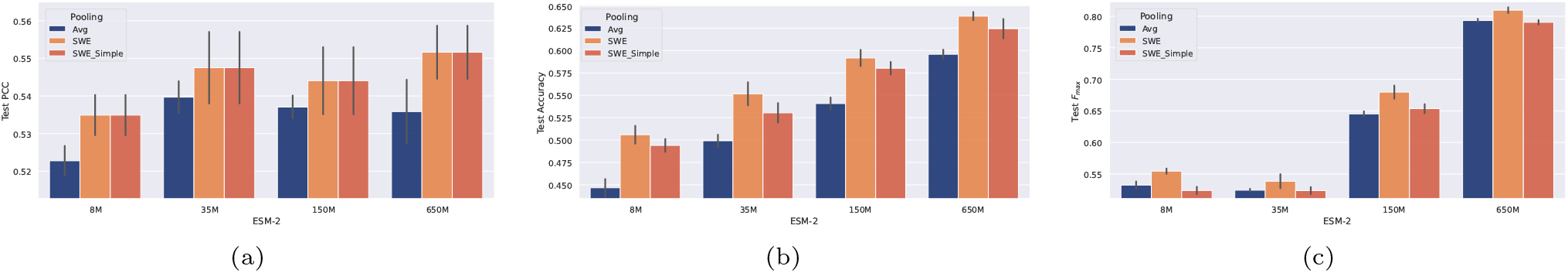
Performance comparison of SWE, SWE_Simple, and average pooling in the (a) drug-target affinity prediction, (b) sub-cellular localization, and (c) EC prediction tasks across four ESM-2 models.

### Sub-Cellular Localization

Figure 4-b demonstrates the performance comparison between SWE, SWE_Simple, and average pooling on the sub-cellular localization task. While both SWE and its simple version outperform average pooling for all four ESM-2 PLMs, we observe a performance gap between SWE and SWE_Simple, implying that training the reference and slicer parameters for this task leads to a higher accuracy than keeping them frozen.

### Enzyme Commission Prediction

Figure 4-c shows the EC prediction performance of SWE and SWE_Simple compared to average pooling across four ESM-2 PLMs. While the simple version of SWE performs on par with average pooling on this task, training SWE’s reference and slicer parameters results in a performance boost across all ESM-2 PLMs, providing further proof of the efficacy of the proposed pooling method.

## Discussion

We introduce a novel approach for protein representation based on optimal transport principles. We anticipate that our method could have applications beyond PLMs to other foundation models in biology (e.g., for DNA sequence models). Our approach summarizes per-token embeddings in terms of their distance from a set of reference embeddings that are learned from the data. Compared to the conventional approach of average pooling, this learnability provides our approach with greater flexibility in capturing task-specific knowledge. Across a fairly diverse array of datasets, we observed that pooling works acceptably well for very large PLMs, where the greater complexity of the sequence representation compensates for the lack of learnability in the summarization layer. However, these large models have memory requirements that are beyond the capabilities of many GPUs. Sophisticated summarization approaches will be crucial in maximizing the power of the smaller models that can fit on common GPUs and could further unlock parameter-efficient fine-tuning opportunities for such pre-trained models [40; 37]. Our proposed method provides a relatively lightweight aggregation technique to improve the performance of PLMs. The additional complexity is especially negligible for the simple version of SWE, which only includes hundreds of learnable parameters, as compared to the millions-billions of parameters included in state-of-the-art PLMs.

Future work could focus on further exploring the interpretability of our method. Potential research directions include explicitly using active/binding sites in a dictionary learning-type setup, where the key residues in the interaction surface inform the selection of reference set elements, and incorporating auxiliary loss functions to encode greater biological intuition into the proposed aggregation operation. Such approaches would enhance the biological relevance of the representations produced by our method. Our work thus opens up new pathways for enhancing the interpretability and efficiency of PLM-based approaches in biology.

## Appendix

### Training Settings and Hyperparameters

For the DTI and SCL tasks, we train the SWE aggregation parameters using the AdamW optimizer [27] with a learning rate of 10^−4^ and cosine annealing schedule with a restart duration of 10 epochs. For the EC task, we train the SWE aggregation parameters using the Adam optimizer [19] with a learning rate of 10^−3^, reducing it by a factor of 0.6 if the performance does not improve after 5 epochs. The batch size is set to 32 for the DTI and SCL tasks and 8 for the EC task.

### Binary DTI Prediction AUROC Results

Figure 5 shows the AUROC comparison between the proposed pooling method and the baselines for the DTI prediction task on DAVIS and Binding-DB for ESM-2 and ProGen2 models. The overall trends and gains of SWE with respect to baseline pooling operations are consistent with our AUPR results in Figure 2.

**Fig. 5.**
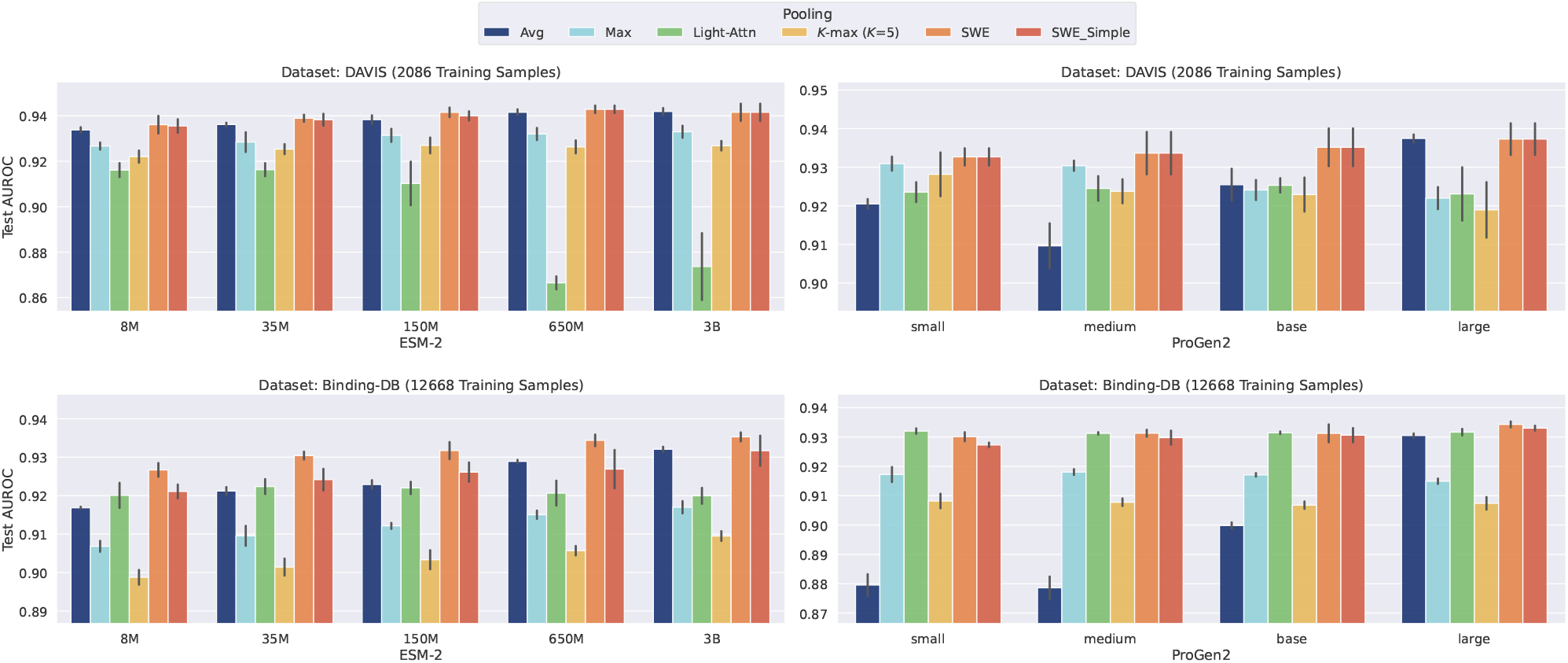
Test AUROC performance of SWE and SWE_Simple as compared to baseline pooling methods for the binary drug-target interaction task on the DAVIS (top) and Binding-DB (bottom) datasets. Performance is evaluated with two families of PLMs, namely ESM-2 (left) and ProGen2 (right).

Moreover, Figure 6 shows the AUROC gains of SWE over average pooling across different protein lengths. Similar to the AUPR gains in Figure 3, SWE’s gains over average pooling increase with the sequence length and decrease with the PLM size.

**Fig. 6.**
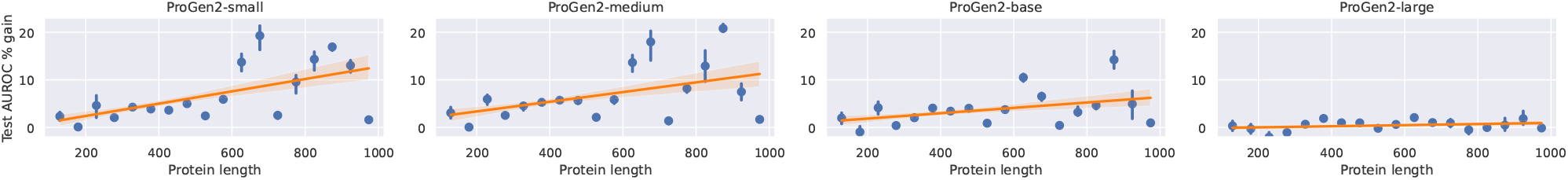
Test AUROC gains of SWE over average pooling vs. the lengths of the proteins’ amino acid sequences across four ProGen2 PLMs on Binding-DB.

### Protein-Protein Interaction (PPI) Prediction

In this task, we focus on predicting whether or not two given proteins will interact with each other. Specifically, we embed each protein’s amino acid sequence using the same PLM and aggregation pipeline in parallel, and then we leverage the cosine similarity between the protein-level representations to estimate their interaction probability. For a given PPI training dataset consisting of *N* (protein, protein, label) triplets 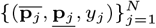, this task seeks to solve the following supervised learning problem:

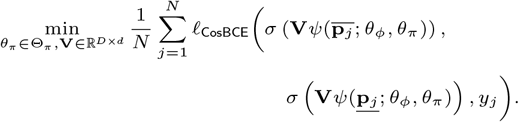

We use the “gold standard” dataset provided in [5] to evaluate our proposed SWE aggregation mechanism in this task, which comprises balanced PPI data without any leakage between training, validation, and testing samples. For this specific task, we consider four options for the number of slices, *L* ∈ {128, 256, 512, 1024}, as well as four options for the number of reference points, *m* ∈ {128, 256, 512, 1024}, leading to 16 different SWE configurations. While the default batch size is 32, for experiments with *L* = 1024 slices, we use a reduced batch size of 24 due to computational limitations. We report the mean and standard deviation of the test/validation performance for the SWE configuration (i.e., (*L, m*) pair) with the highest validation AUPR across the 50 training epochs.

Table 1 compares the validation and test performance of our proposed SWE aggregation method with average pooling across the considered ESM-2 PLMs. As the table shows, SWE generally outperforms average pooling in terms of F1-score and recall, while performing on par with average pooling in terms of accuracy, and underperforming for the other metrics, including precision and specificity. These results are consistent with the ones reported by [40], where more expressive models and heavier fine-tuning lead to superior F1 and recall levels, while lowering the other metrics. Further research, including using optimal partial transport-(OPT-)based embedding methods [2; 3], is required to strike the right balance among the different metrics through which PPI prediction quality is measured.

**Table 1.** PPI validation and test results on the “gold standard” dataset in [5] for the SWE and average pooling methods across four different ESM-2 PLM backbones. Following [40], for a comprehensive evaluation, we report the performance using seven different metrics, including accuracy, F1-score, Matthews correlation coefficient (MCC), AUPR, precision, recall, and specificity, with mean and standard deviation across five different random seeds. Underlined numbers indicate the best aggregation performer in each metric for each phase (validation/test) and each PLM. Bold numbers indicate the best performer in each metric for each phase (validation/test).

| ESM-2 PLM |  | Aggregation | Accuracy | F1 | MCC | AUPR | Precision | Recall | Specificity |
| --- | --- | --- | --- | --- | --- | --- | --- | --- | --- |
| Validation | 8M | SWE | 0.607 ± 0.003 | 0.626 ± 0.008 | 0.216 ± 0.006 | 0.652 ± 0.004 | 0.619 ± 0.006 | 0.671 ± 0.023 | 0.701 ± 0.013 |
|  |  | Avg | 0.606 ± 0.002 | 0.614 ± 0.004 | 0.216 ± 0.004 | 0.653 ± 0.002 | 0.637 ± 0.004 | 0.643 ± 0.012 | 0.731 ± 0.008 |
|  | 35M | SWE | 0.617 ± 0.002 | 0.632 ± 0.009 | 0.235 ± 0.004 | 0.669 ± 0.003 | 0.627 ± 0.002 | 0.693 ± 0.024 | 0.690 ± 0.009 |
|  |  | Avg | 0.617 ± 0.001 | 0.628 ± 0.008 | 0.237 ± 0.002 | 0.667 ± 0.001 | 0.648 ± 0.001 | 0.666 ± 0.024 | 0.735 ± 0.005 |
|  | 150M | SWE | 0.623 ± 0.002 | 0.633 ± 0.006 | 0.248 ± 0.003 | 0.677 ± 0.003 | 0.633 ± 0.005 | 0.663 ± 0.021 | 0.681 ± 0.011 |
|  |  | Avg | 0.624 ± 0.002 | 0.635 ± 0.006 | 0.251 ± 0.004 | 0.679 ± 0.003 | **0.661 ± 0.004** | 0.676 ± 0.023 | **0.754 ± 0.008** |
|  | 650M | SWE | **0.629 ± 0.002** | **0.668 ± 0.001** | **0.259 ± 0.004** | **0.685 ± 0.002** | 0.630 ± 0.008 | **0.810 ± 0.004** | 0.643 ± 0.023 |
|  |  | Avg | 0.626 ± 0.001 | 0.650 ± 0.004 | 0.251 ± 0.002 | 0.684 ± 0.002 | 0.651 ± 0.003 | 0.732 ± 0.030 | 0.732 ± 0.008 |
| Test | 8M | SWE | 0.637 ± 0.004 | 0.638 ± 0.013 | 0.274 ± 0.008 | 0.688 ± 0.005 | 0.636 ± 0.007 | 0.642 ± 0.030 | 0.632 ± 0.027 |
|  |  | Avg | 0.639 ± 0.000 | 0.637 ± 0.005 | 0.279 ± 0.001 | 0.690 ± 0.000 | 0.641 ± 0.004 | 0.633 ± 0.013 | **0.645 ± 0.013** |
|  | 35M | SWE | 0.646 ± 0.003 | 0.661 ± 0.010 | 0.294 ± 0.004 | 0.696 ± 0.005 | 0.635 ± 0.011 | 0.690 ± 0.035 | 0.602 ± 0.040 |
|  |  | Avg | 0.648 ± 0.000 | 0.651 ± 0.010 | 0.297 ± 0.001 | 0.699 ± 0.000 | 0.646 ± 0.008 | 0.658 ± 0.029 | 0.639 ± 0.028 |
|  | 150M | SWE | 0.649 ± 0.002 | 0.664 ± 0.007 | 0.299 ± 0.005 | 0.703 ± 0.002 | 0.636 ± 0.005 | 0.696 ± 0.020 | 0.601 ± 0.020 |
|  |  | Avg | 0.651 ± 0.001 | 0.654 ± 0.009 | 0.303 ± 0.003 | 0.708 ± 0.002 | **0.649 ± 0.006** | 0.659 ± 0.025 | 0.643 ± 0.023 |
|  | 650M | SWE | 0.650 ± 0.005 | **0.683 ± 0.007** | 0.307 ± 0.007 | 0.705 ± 0.005 | 0.625 ± 0.012 | **0.754 ± 0.033** | 0.545 ± 0.042 |
|  |  | Avg | **0.657 ± 0.002** | 0.670 ± 0.006 | **0.315 ± 0.003** | **0.717 ± 0.001** | 0.646 ± 0.007 | 0.696 ± 0.022 | 0.618 ± 0.024 |

#### Interpretability of the Learned SWE Representations

One of the desirable properties of the proposed embeddings is that the sliced-Wasserstein distance of the distributions underlying the token-level embeddings of two given protein sequences can be approximated by the average distance of their Monge couplings to the reference across different slices [31]. In particular, for two proteins 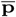 and **<u>p</u>** with Monge coupling matrices 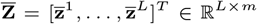 and 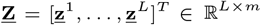, respectively, we can approximate their pairwise sliced-Wasserstein distance as

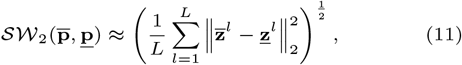

where, with a slight abuse of notation, we use 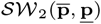 to denote the sliced-Wasserstein distance between token-level representations of 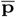 and **<u>p</u>** at the output of the PLM backbone.

The approximation in (11) allows us to visualize the pairwise distance of interacting and non-interacting proteins in the embedding space. Figure 7 shows the distributions of (approximate) SW distances between proteins in the training dataset separated by whether or not they interact, where the embeddings are generated by the 650M-parameter ESM-2 backbone and a trained SWE aggregation module with *m* = 1024 reference points and *L* = 1024 slices. As the figure demonstrates, there is a separation between the two histograms, with interacting proteins landing closer to each other in the embedding space from a sliced-Wasserstein distance point of view as compared to non-interacting pairs of proteins.

**Fig. 7.**
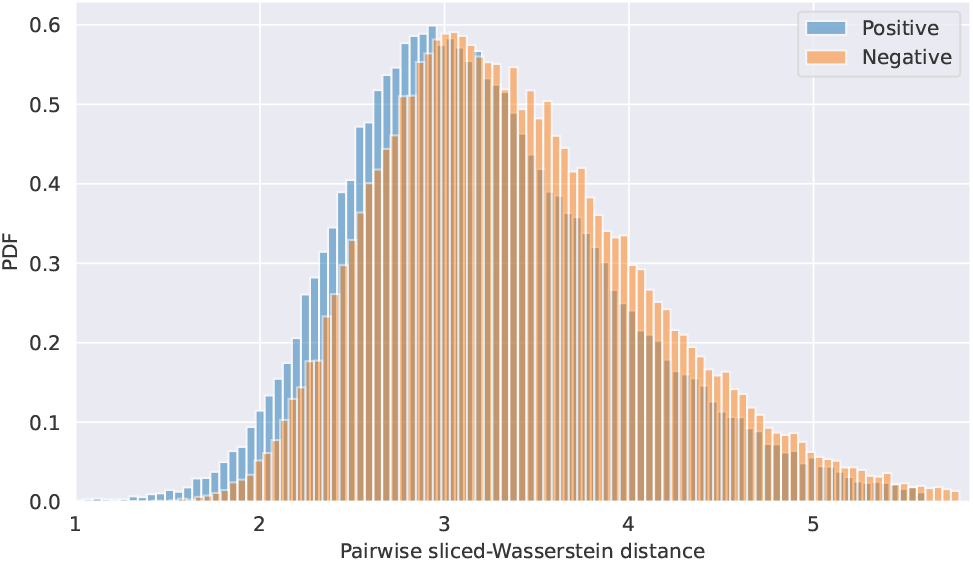
Histograms of sliced-Wasserstein distance between embeddings of interacting and non-interacting proteins at the output space of a pre-trained ESM-2 PLM with 650M parameters. The SW distance is calculated using a trained SWE pooling module with *m* = 1024 reference points and *L* = 1024 slices.

## Footnotes

1 https://tdcommons.ai/benchmark/dti_dg_group/BindingDB_Patent

## Notes

### Competing Interest Statement

The authors have declared no competing interest.

### Summary of Updates

Key changes compared to the previous version of the manuscript include the following: - Improving the performance of the proposed method by adaptively keeping the aggregation parameters frozen based on the training set size and the protein language model (PLM) architecture. - Introducing a parameter-efficient version of our approach, called “SWE_Simple,” which in many cases outperforms average pooling. - Adding evaluations on larger ESM-2 PLMs, as well as an array of ProGen2 PLMs. - Evaluating SWE on additional downstream prediction tasks, including subcellular localization and enzyme commission prediction. - Including more baseline pooling operations to paint a more comprehensive picture of SWE's performance vs. alternative methods. - Demonstrating that the gains of SWE over average pooling are most pronounced for longer protein sequences.

